# The median and the mode as robust meta-analysis methods in the presence of small study effects

**DOI:** 10.1101/288050

**Authors:** Fernando Pires Hartwig, George Davey Smith, Amand Floriaan Schmidt, Jack Bowden

## Abstract

Meta-analyses based on systematic literature reviews are commonly used to obtain a quantitative summary of the available evidence on a given topic. Despite its attractive simplicity, and its established position at the summit of the evidence-based medicine hierarchy, the reliability of any meta-analysis is largely constrained by the quality of its constituent studies. One major limitation is small study effects, whose presence can often easily be detected, but not so easily adjusted for. Here, robust methods of estimation based on the median and mode are proposed as tools to increase the reliability of findings in a meta-analysis. By re-examining data from published meta-analyses, and by conducting a detailed simulation study, we show that these two simple methods offer notable robustness to a range of plausible bias mechanisms, without making any explicit modelling assumptions. In conclusion, when performing a meta-analysis with suspected small study effects, we recommend reporting the mean, median and modal pooled estimates as a simple but informative sensitivity analyses.

## 1. Introduction

Meta-analysis is a statistical technique for obtaining a quantitative summary of the totality of evidence on a given topic, as part of a broader systematic review.^1,2^ In its archetypal form, it provides an overall effect estimate for a well-defined intervention that has been assessed across several independent studies. In addition, meta-analyses provide a further opportunity to explore between study heterogeneity, which might highlight novel patient subgroups with contrasting treatment responses.^1,2^

Unfortunately, between-study heterogeneity may also indicate the presence of bias, which wish to understand, but ultimately remove, from the final analysis. For example, it is generally accepted that results in agreement with the prevailing wisdom and achieving conventional levels of significance are more likely to be reported and published than negative or inconclusive findings. Therefore, published studies are more likely to present large effect estimates, corresponding to a biased sample of all studies.^2^ Larger studies are affected to a lesser extent than smaller studies, most obviously because an increased sample size raises the likelihood of achieving conventional statistical significance when the true effect to be measured is non-zero. Large studies also require a sizeable financial outlay from funding agencies and often represent the collective effort of many cross-institutional researchers. These factors provide added impetus to place the results in the public domain, regardless of their conclusions. Moreover, small studies are more often early-phase trials (which may use less stringent designs) than larger studies, or may preferentially include high risk patients in order to improve power. This phenomenon, which encompasses many different mechanisms, is referred to generically as “small study effects”.^2^

It is often difficult to identify whether any observed correlation between study size and reported treatment effect is due to “true” between-study differences, selective reporting and publication, or a combination of both.^2^ Many different methods to detect and/or correct for small study effects have been proposed. One of the earliest of such methods is the funnel plot (where study-specific point estimates are plotted against their precision), which has been proposed more than 30 years ago. Asymmetry in the funnel plot may be indicative of selective reporting and publication, although “true” between-study differences correlated with precision can exist.^3^ This motivated the development of methods that “correct” for asymmetry, such as Egger regression and trim-and-fill.^3–5^ However, because these methods make either implicit or explicit assumptions about the selection process, their performance suffers acutely when the true bias mechanism is different. This can easily result in a bias-adjusted estimate that is further from the truth than the original result. This motivates our proposal of two simple methods which are naturally robust to small study effects, whilst making no assumptions about its precise nature.

## 2. Methods

Before discussing our proposed estimation procedures in detail, we first describe the underlying data generating model they will be evaluated against. This slightly unusual step is necessary for the reader to understand when each method can, in theory, identify the true treatment effect.

### 2.1. Data generating model

We start by defining a summary data generating model with *K* studies indexed by *j*(*j* = 1,2, …, *K*). Each study reports an estimated mean difference between randomised groups (e.g., one to an experimental intervention and one to standard intervention) denoted by 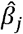 where:

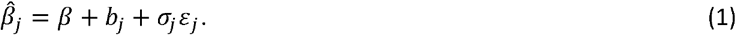

Here:

1. *β* is the true effect of the exposure on the outcome;
2. *b*_*j*_ denotes the bias/heterogeneity parameter for study *j*;
3. *σ*_*j*_ is the standard error of the mean difference;
4. *ε*_*j*_~*N*(0,1,*l*_*j*_,*u*_*j*_) is a draw from a standard truncated Normal distribution with lower limit *l* and upper limit *u*. When *l* = −∞ and *u* = ∞, then *ε*_*j*_ denotes pure random error due to sampling variation.
5. The parameters *b*_*j*_, *σ*_*j*_, *l*_*j*_, and *u*_*j*_ are all allowed to depend on the study size, *n*_*j*_.

We will use *b*_*j*_ and *ε*_*j*_ to variously induce heterogeneity, effect modification, and small study bias in the data, as described below.

### 2.2. General principles of our bias model

We will explore two types of small study bias: bias due to systematic differences between small and large studies due to study quality (type (a)), and bias due to the specific environment of selective reporting and publication in operation at the time when study *j* was conducted (type (b)).

For type (a), we imagine that bias is a fundamental property of each study, in that the true treatment effect for study *j* is *β* + *b*_*j*_, where *b*_*j*_ is a simple, non-increasing function of study size (*n*_*j*_). That is:
*b*_*j*_ ≤ *b*_*k*_, whenever *n*_*k*_ ≤ *n*_*j*_.

Without loss of generality we assume that the bias is always positive, so that *b*_*j*_ ≥ 0. We will investigate cases where the bias disappears only asymptotically as a study size grows infinitely large, and cases where the bias disappears beyond a threshold study size, *N*. That is:
*b*_*j*_ → 0 as *n*_*j*_ → ∞, or *b*_*j*_ = 0 if *n*_*j*_ ≥ *N* for some (large) *N*.

Type (b) bias is not a fundamental component of the study itself, but instead the result of selective reporting and publication (i.e., dissemination bias). We induce this through the random error component of model (1), *ε*_*j*_, in the following manner.

Again, we assume that bias is always positive, so that *E*[*ε*_*j*_|*n*_*j*_] ≥ 0. This corresponds to a situation where the selection process favours the publication of studies that reported positive effect estimates. We achieve this by defining the lower limit of 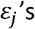 truncated normal distribution, *l*_*j*_, as a non-increasing function of *n*_*j*_. That is:

*l*_*j*_ ≤ *l*_*k*_, and therefore *E*[*ε*_*j*_|*n*_*j*_] ≤ *E*[*ε*_*k*_|*n*_*k*_], whenever *n*_*k*_ ≤ *n*_*j*_.

Similarly to the type (a) bias model, we will explore cases where:

*l*_*j*_ → −∞ and *E*[*ε*_*j*_|*n*_*j*_] → 0 as *n*_*j*_ →∞, or

*l*_*j*_ = −∞ and *E*[*ε*_*j*_|*n*_*j*_] = 0 if *n*_*j*_ ≥ *N* for some large *N*.

A general expression for the expected value of study *j*’s effect estimate 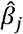, based on *n*_*j*_ participants and in the presence of type (a) and type (b) bias, is therefore:

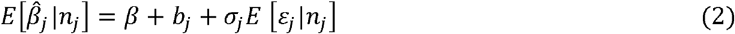

An important distinction between type (a) and type (b) bias is their respective effect on the variance of the study-specific estimates. Type (a) bias will generally increase their variability, leading to over-dispersion, or heterogeneity. Type (b) bias, by contrast, can have the opposite effect of reducing their variability, because of the truncation in the distribution of *ε*_*j*_. That is, in the presence of this bias, Var[*ε*_*j*_|*n*_*j*_] will generally be less than 1, and Var[*ε*_*j*_|*n*_*j*_] ≥ Var[*ε*_*k*_|*n*_*k*_] whenever *n*_*k*_ ≤ *n*_*i*_. This phenomenon leads to under-dispersion across the set of study-specific estimates constituting the meta-analysis.

### 2.3. Robust central tendency statistics in meta-analysis

We now introduce three estimators for the overall effect *β*: the standard approach plus two novel approaches, and discuss their ability to return consistent estimates for data generated under model (1). For the purposes of clarity only, we will momentarily assume that *b*_*j*_ is the sole source of bias in equation (1) - i.e., that *E*[*ε*_*j*_|*n*_*j*_] = 0.

#### 2.3.1. The weighted mean

A standard fixed effect meta-analysis would estimate the effect size parameter *β* as an inverse-variance weighted average (or pooled mean) of the individual study estimates. That is:

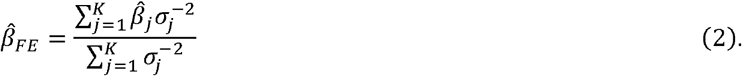

However, if even a single study contributes a biased estimate to the meta-analysis (e.g., via a non-zero *b*_*j*_), then the pooled mean will also generally be biased. That is, using the notation of formula (1):

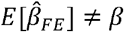 in general, whenever *b*_*j*_ > 0 for some study *j* in 1,…, *K*.

For this reason, in the language of robust statistics, the mean is said to have a 0% “breakdown” level.

#### 2.3.2. The weighted median

The weighted median^6^ estimate is defined as the 50th percentile of the inverse-variance weighted empirical distribution of the study specific estimates, which can be calculated as follows. Assume that the 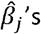 are sorted in ascending order, so that 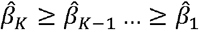. Let the standardised inverse-variance weights for study *j* be defined as 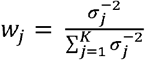 and sort them in the same order as the 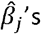. Finally, let 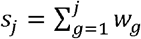 denote the sum of standardised weights up to and including the *j*th study. This means that 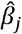 is the 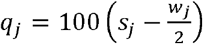 th percentile of the weighted empirical distribution of 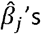.

The weighted median estimate is the 50% percentile of this distribution, so it will be equal to 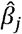 if *S*_*j*_ = 0.5. In general, no study lies exactly at the 50^th^ percentile, so this quantity is estimated in practice by linear interpolation between its neighbouring estimates 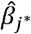 and 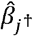, which correspond to the effect estimates reported by the studies located immediately before and after the 50% percentile, respectively (i.e., 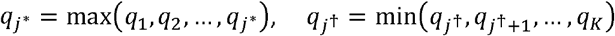, and 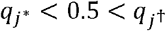). In this case, the weighted median estimate 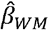 is:

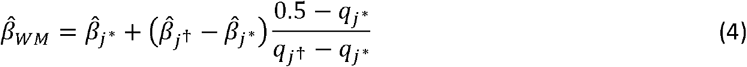

The weighted median does not require that all 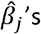 are consistent estimates for the true effect *β*. More specifically, 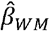 is consistent if up to (but not including) 50% of the total weight in the analysis comes from biased studies - i.e., 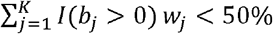. This means that the weighted median has a breakdown level of 50%.

#### 2.3.3 The mode-based estimate

The mode-based estimate^7^ (MBE) works by exploiting the Zero Modal Bias Assumption (ZEMBA), which requires that the most common value of the bias parameter *b*_*j*_ is zero. If ZEMBA holds, the mode of all 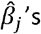 (hereafter referred to as 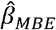) is a consistent estimate of the true effect *β*, even if the majority of 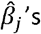 are biased.

More formally, 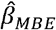 consistent if *w*_0_ > ma*x*(*w*_1_,*w*_2_, …, *w*_*v*_), where (in a departure from the convention introduced for the weighted median) *w*_0_ now denotes the sum of weights provided by studies with zero bias, and *w*_1_ *w*_2_ and *w*_*v*_ are the sum of weights provided by studies that have the smallest, the second smallest and the largest identical bias terms, respectively.

It is possible to define many pooled effect estimators that exploit ZEMBA in different ways. Here, as in Hartwig et. al,^7^ we used the mode of the smoothed, inverse-variance weighted empirical density function of all 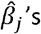 as the MBE. More specifically, 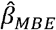 the value of *x* that maximizes *f*(*x*) (i.e., 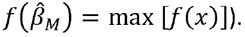. *f*(*x*) is the normal kernel density function:

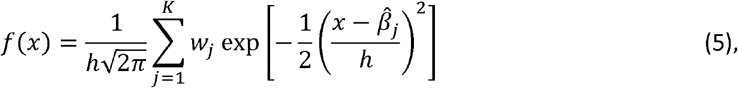

where *h* is the smoothing bandwidth parameter.^8^ This parameter regulates a bias-variance trade-off, with smaller values of *h* reducing both bias and precision. We used the modified Silverman’s bandwidth selection rule proposed by Bickel et al.^9^

The exact breakdown level of the MBE depends on max(*w*_1_,*w*_2_, …,*w*_*v*_), which is unknown. If all biased studies estimate the exact same effect parameter, then ZEMBA will only be satisfied if up to (but not including) 50% of the weights comes from biased studies. The upper limit of the breakdown level is up to (but not including) 100%, and corresponds to the situation where all invalid studies estimate different effect parameters. Therefore, the breakdown level of the MBE ranges from 50% to 100%.

The weighted median and MBE were originally proposed as robust tools for summary data Mendelian randomization,^6,7^ which is analogous to a meta-analysis.

### 2.4. Illustrating the identifying assumptions of the mean, median and mode

Figure 1 illustrates the assumptions underlying the pooled mean, median and mode in a hypothetical meta-analysis of 10 studies, sorted in ascending order of their 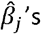. The true effect *β* is zero. For simplicity, all studies have the same weight and no sources of heterogeneity other than bias are present. Chiefly:

**Figure 1.**
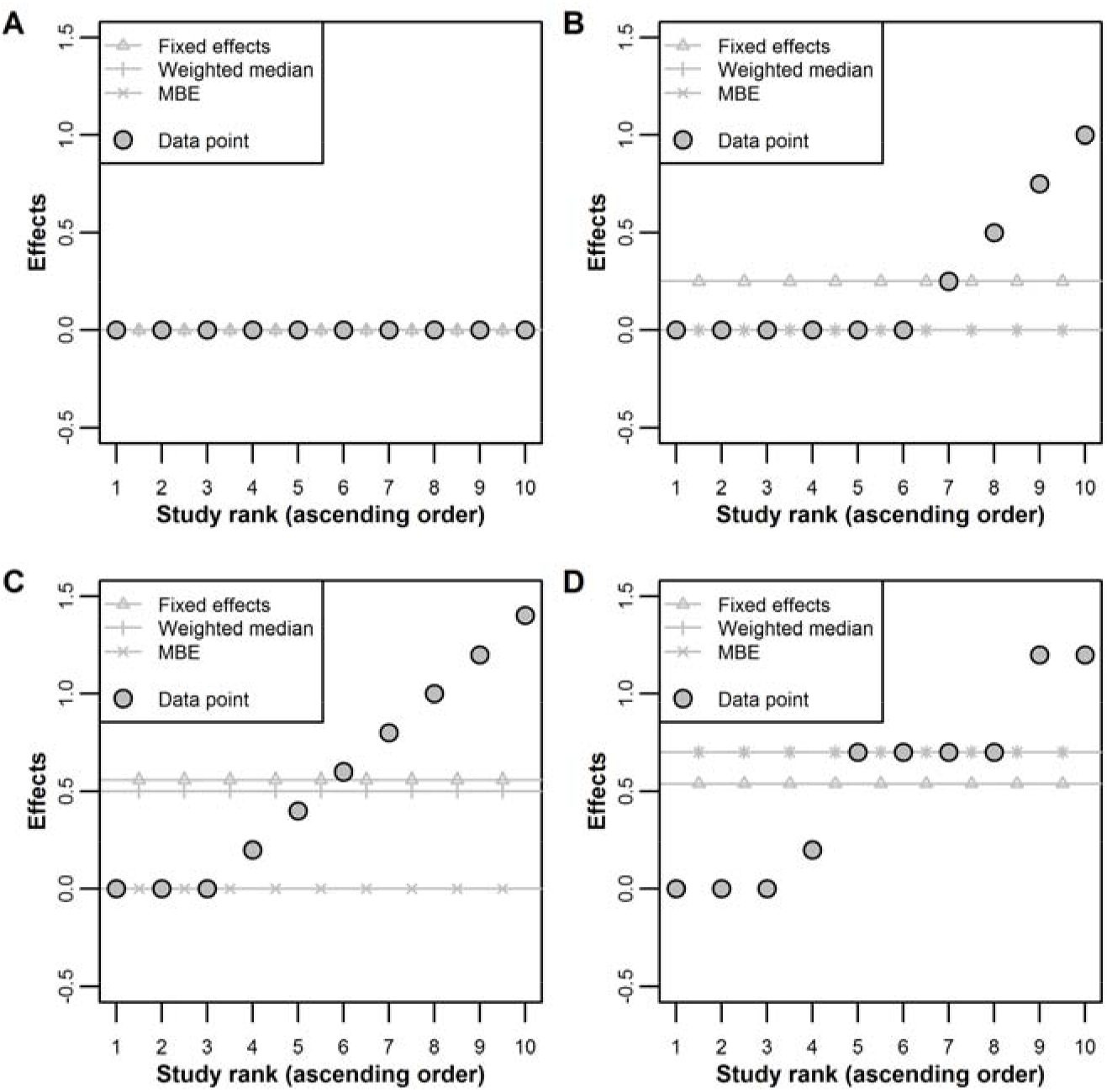
Illustration of the assumptions underlying the weighted median and the mode-based estimate (MBE) methods. Studies are assumed to have the same weights in the meta-analysis, and are sorted in ascending order of point estimate. The true effect is zero. A: no heterogeneity between studies. B: 4 out of 10 studies are biased. C: 7 out of 10 studies are biased, but unbiased studies comprise the largest subgroup of studies that reported the same result, all biased studies reported different effects. D: 7 out of 10 studies are biased, and biased studies comprise the largest subgroup of studies that reported the same result.

- When all 10 studies (i.e., 100%) are unbiased (Panel A), all three methods identify the true effect (zero);
- When 4 out of 10 studies are biased (Panel B), or whenever less than 50% of studies are biased in general, the mean is biased, but the median and the mode are unbiased;
- When 7 out of 10 studies are biased (Panel C), or whenever more than 50% of studies are biased in general, and ZEMBA is satisfied, both the mean and the median are biased, but not the mode;
- When more than 50% of the studies are biased (Panel D) and ZEMBA is violated, all methods are biased.

One attractive property of the weighted median and MBE is that they are naturally robust central tendency statistics, and do not make any specific assumptions about the selection mechanism at play. Therefore, they might be robust to a range of reasonable small study effects models. However, as Figure 1 illustrates, these methods are not guaranteed to provide consistent estimates of *β*, failing to do so when their identifying assumptions are violated. Nevertheless, these assumptions are weaker than the assumptions required by the standard pooled mean.

### 2.5. Simulation study

We performed a simulation study to evaluate the performance of different meta-analysis methods in a range of small study effects models. We evaluated the following methods: fixed effects model, Egger regression, trim-and-fill, weighted median and MBE.

Summary data were generated using equation (1). We assume that each study measured a binary exposure variable *X*~Bernoulli(0.5) (e.g., an intervention: yes=1, no=0) and a continuous outcome variable *Y* with variance equal to one. Therefore, the pooled standard error of the mean difference is one for all values of *j*, and 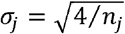, where *n*_*j*_ is the sample size of the *j*th study. We will assume studies range in size from *n*_1_ to *n*_2_ uniformly, so that *n*_*j*_~Uniform(*n*_1_,*n*_2_).

#### 2.5.1. Type (a) bias

The value of the bias term *b*_*j*_ was defined as the following linear function of study size: 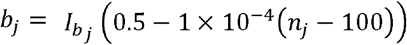. From this model, if *n*_*j*_ = 100 (the smallest study size in our simulations), then 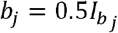. If *n*_*j*_ = 5000 (the largest study size in our simulations), then 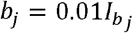.

The indicator function 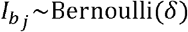, with *δ* ∈ [0,1], dictates the presence 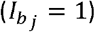 or absence 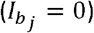 of bias. Therefore, the expected number of studies suffering from bias equals *δK*.

#### 2.5.2. Type (b) bias

As described above, this bias was generated through *ε*_*j*_, by varying *l*_*j*_ according to study size. Typically, dissemination bias models assume that results that achieve conventional levels of statistical significance are more likely to be published. Therefore, in our simulations, *l*_*j*_ was defined to correspond the maximum one-sided P-value (null hypothesis: true mean difference ≤0) allowed for publication for a given study size (*p*_*j*_). That is:

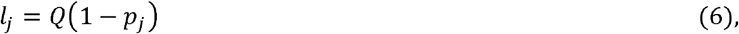

where *Q(p)* is the quantile function for the Student’s t distribution with *n*_*j*_ − 1 degrees of freedom. For example, if *p*_*j*_ = 0.025 and *n*_*j*_ = 1000, then *l*_*j*_ = *Q*(1 − *p*_*j*_) = *Q*(0.975) ≈ 1.96. This situation can be interpreted as studies with 1000 participants only being publishable if the reported one-sided P-value is ≤ 0.025.

We generated dissemination bias by defining *p*_*j*_ as various functions of *n*_*j*_, as described below,

i. *p*_*j*_ as a continuous function of *n*_*j*_(up to *N*). In this model, 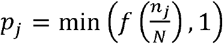, where 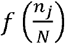 is some non-decreasing function of 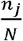 and *N* is some upper threshold study size threshold at which *p*_*j*_ = 1 (and therefore *l*_*j*_ = −∞) for all *n*_*j*_ ≥ *N*. We compared three distinct functions:
  - Identity: 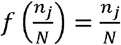
  - Square root: 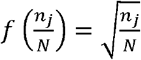 and
  - Quadratic: 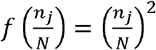.
ii. *p*_*j*_ as a step function of *n*_*j*_. In this model, *p*_*j*_ is defined as a piecewise function of *n*_*j*_ classifying studies into small, medium or large. That is:

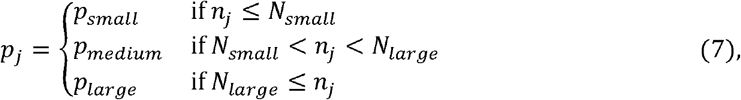

where 1 ≤ *N*_*small*_ ≤ *N*_*large*_ and 0 ≤ *p*_*small*_ ≤ *p*_*medium*_ ≤ *p*_*large*_ ≤ 1. Therefore, studies classified in the same group have the same P-value requirements for publication, and the relationship between *p*_*j*_ and *n*_*j*_ follows a step function. The relationship between *p*_*j*_ and *n*_*j*_ in each one of these four models is illustrated in Supplementary Figure 1.

#### 2.5.3. Simulation scenarios

We evaluated the meta-analysis methods described above in the simulation scenarios described below. In all cases, *K* was set to 5,10, 30 or 50.

1. Scenario 1: the true causal effect was zero (i.e., *β* = 0), as in scenarios 2-6, and no small study effects. Therefore, equation (1) simplifies to 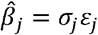 where *ε*_*j*_~*N*(0,1, − ∞,∞). Study size varied as follows: i) *n*_1_ = 100, *n*_2_ = 1000; and ii) *n*_1_ = 1000, *n*_2_ = 5000.
2. Scenario 2: type (a) bias only, yielding the data generating model 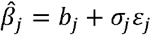 for *ε*_*j*_~*N*(0,1, − ∞, ∞). Study sizes ranged between *n*_1_ = 100 and *n_2_ =* 5000 (these values were also used in scenarios 3-6), and the proportion of biased studies *δ* was varied between 0 and 1 in steps of 0.1.
3. Scenarios 3-5: Type (b) bias only, yielding the data generating model 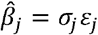, *ε*_*j*_ |*n*_*j*_ ~ *N*(0,1,*l*_*j*_, ∞), and 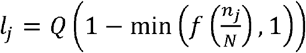 We assumed a linear (scenario 3), square root (scenario 4) or quadratic (scenario 5) relationship between *p*_*j*_ and *n*_*j*_. *N* was set to 1 (i.e., no small study effects), 1500, 3000, 4500 and 6000.
4. Scenario 6: type (b) bias only, assuming a step-function relationship between *p*_*j*_ and *n*_*j*_. This used the same model as in Scenarios 3-5 except with *p*_*j*_ defined following equation (7) and where *P*_*small*_ = 0.025, *P*_*medium*_ = 0.15 and *P*_*large*_ = 1 (the latter implying that large studies have no P-value requirements for publication) were kept constant. Cut-offs to classify studies into small, medium or large varied as follows: i) *N*_*small*_ = *N*_*large*_ = 1 (i.e., no small study effects); ii) *N*_*small*_ = 500, and *N*_*large*_ = 1000; iii) *N*_*small*_ = 1000, and *N*_*large*_ = 2000; iii) *N*_*small*_ = 2000, and *N*_*large*_ = 4000.
5. Scenario 7: Identical to scenario 1, except with *β* = 0.02.

The functional relationship between the bias (i.e., 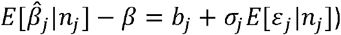 and *n*_*j*_, and between standard error (i.e., 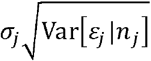 and *n*_*j*_ for each one of the above scenarios is illustrated in Figure 2.

**Figure 2.**
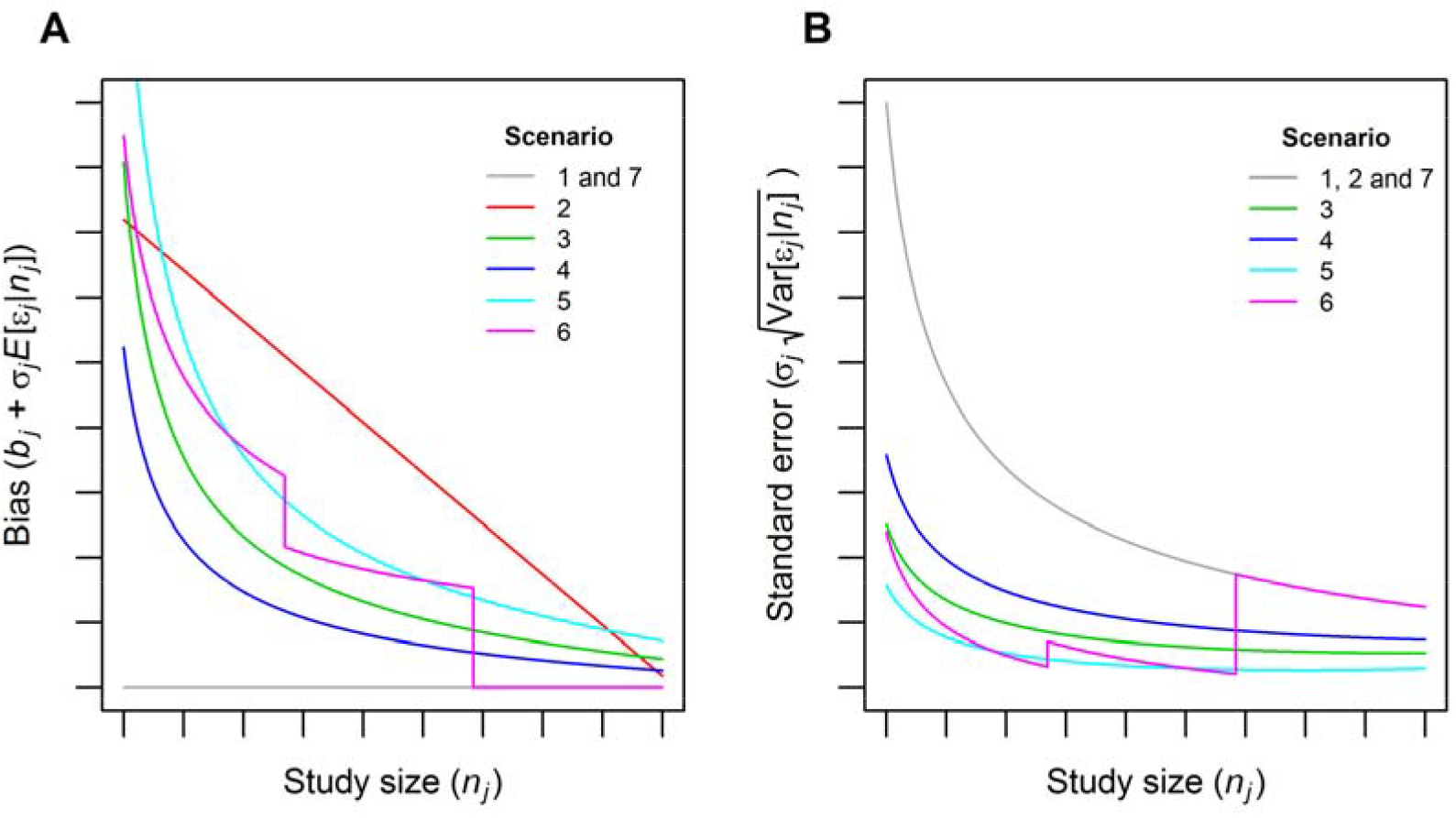
Illustration of the relationship between bias and *n*_*j*_ (panel A), and between standard error and *n*_*j*_ (panel B), induced by different models of small study effects.

#### 2.5.4. Statistical analysis

In the simulation analysis, mean pooled effect estimates, standard errors, coverage and power of 95% confidence intervals were computed for the weighted mean, weighted median, MBE, Egger regression and trim-and-fill methods across 10,000 simulated datasets. Standard errors of the weighted median and the MBE are calculated using parametric bootstrap, which naturally incorporates any between-study heterogeneity into their confidence intervals.

All analyses were performed using R (www.r-proiect.org).

### 2.6. Applied examples

We further evaluated our proposed methods and illustrate their application by re-analysing three meta-analysis datasets:

- Catheter dataset: This meta-analysis, originally conducted by Veenstra et al.^10^ evaluated 11 trials comparing chlorhexidine-silver sulfadiazine-impregnated vs. non-impregnated catheters with regards to risk of catheter-related bloodstream infection. These data presented a large correlation between effect estimates and their precision (*r*=0.76 [P-value=0.007]) (which translates into substantial asymmetry on the funnel plot), and high between-study heterogeneity (*I*^2^=60%).
- Aspirin dataset: this meta-analysis, originally conducted by Edwards et al.,^11^ evaluated 63 trials investigating the effect of a single dose of oral aspirin on pain relief (50% reduction in pain), *r* was also strong in magnitude (*r*=-0.70 [P-value=1.6×10^-6^]), but there was low between-study heterogeneity (*I*^2^=10%).
- Streptokinase dataset: meta-analysis originally conducted by Yusuf et al.^12^ and updated by Egger et al.^3^ of 21 trials evaluating the effect of streptokinase therapy on mortality risk. These data presented moderate heterogeneity (*I*^2^=34%), but very little evidence of asymmetry (*r*=0.08, P-value=0.743). These data were used as a positive control, where all methods are expected to give similar answers.

## 3. Results

### 3.1. Simulation study

Simulation scenario 1 indicated that the confidence intervals of the fixed effects, weighted median and MBE are valid in the sense that they all achieve at least 95% coverage under the null (i.e., *β* = 0) and in absence of small study effects, although only the fixed effects method had exact 95% coverage (Supplementary Table 1). Egger regression presented under-coverage when the number of studies was small, but this attenuated as the number of studies increased. Conversely, the trim-and-fill method presented under-coverage that increased with number of studies, indicating that its confidence intervals are invalid (at least in our implementation of the method). The fixed effects method presented the smallest standard errors, followed by the trim-and-fill, which was slightly more precise than the weighted median. The MBE was less precise than the latter, but substantially more precise than Egger regression.

Supplementary Table 2 shows that scenario 2 lead to high values of *I*^2^ and *r*. Under this small study effects model, the weighted median was less biased than the fixed effects model, and the MBE was the least biased among all methods (Figure 3). Those differences became more apparent as the number of studies increased. The trim-and-fill was more biased than the standard fixed effects model, and Egger regression substantially overcorrected.

**Figure 3.**
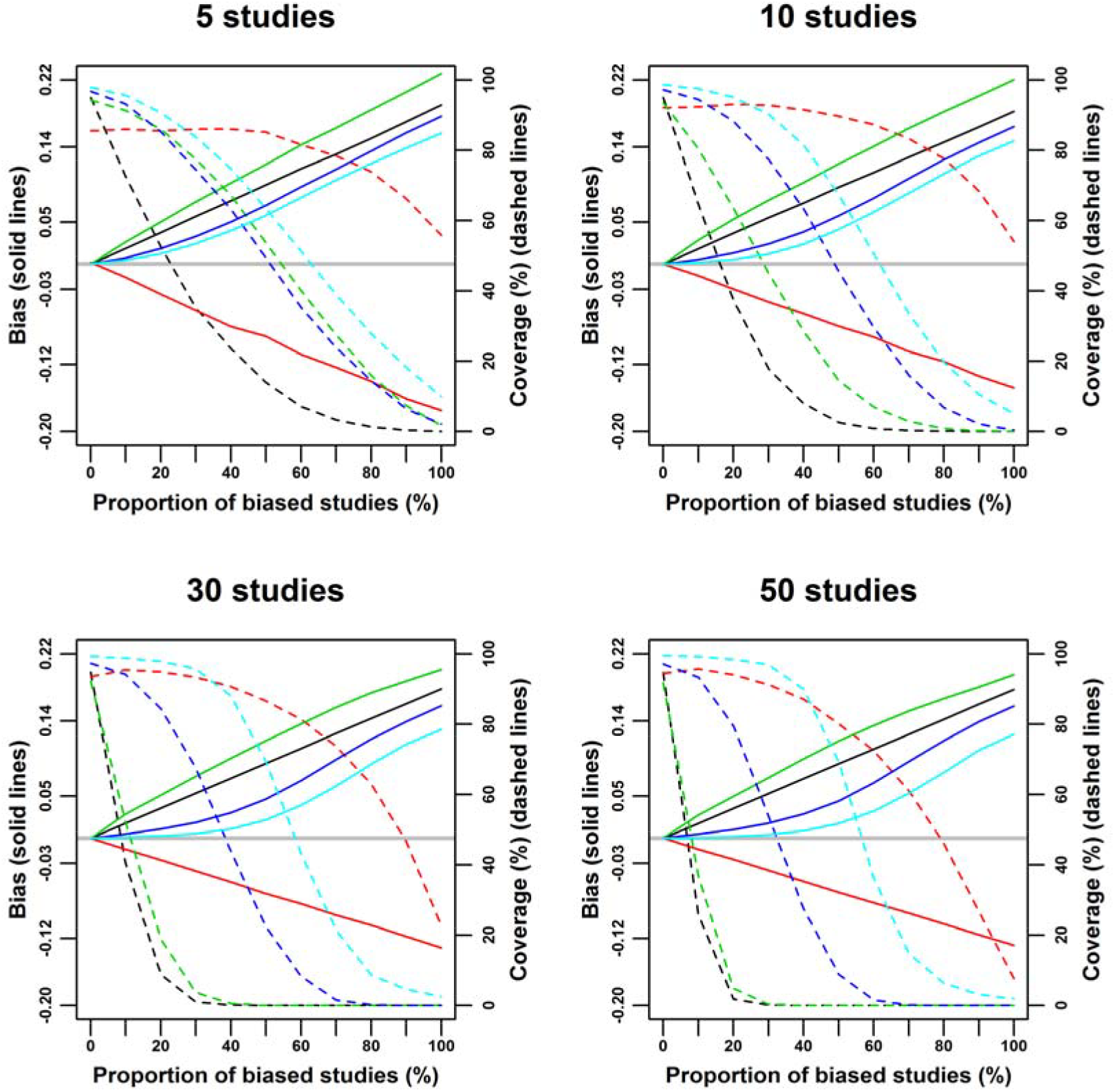
Bias (solid lines) and coverage (dashed lines) of the fixed effects (black), Egger regression (red), trim-and-fill (green), weighted median (dark blue) and mode-based estimate (light blue) under scenario 2: zero true effect (i.e., *β* = 0), small study effects through the bias term *b*_*j*_, and study sizes uniformly ranging from 100 to 5000 individuals. The grey line indicates zero bias.

Scenario 3 lead to high asymmetry, but did not substantially inflate *I*^2^ (Supplementary Table 3), and the bias in the pooled estimates was much smaller compared to scenario 2. Again, Egger regression substantially overcorrected for small study effects, and the weighted median and MBE were less biased than the fixed effects model (Figure 4). However, the performance of the trim-and-fill relative to the weighted median and the MBE was substantially different than in scenario 2: here, it the number of studies is low (*K* = 5), the trim-and-fill performed similarly to the weighted median, but was more biased than the MBE; for *K* = 5, it outperformed the weighted median and performed similarly to the MBE; for larger values of *K*, the trim-and-fill was generally less biased than the other methods, unless all studies were affected by small study effects (in this case, *N* = 6000). However, as the number of studies increased, the trim-and-fill overcorrected for small study effects when *N* = 1500. In general, the differences between the weighted median, the MBE and trim-and-fill were much less marked in this scenario than in scenario 2; indeed, in scenario 3, the coverage of the weighted median and the trim-and-fill was similar for all values of *K*.

**Figure 4.**
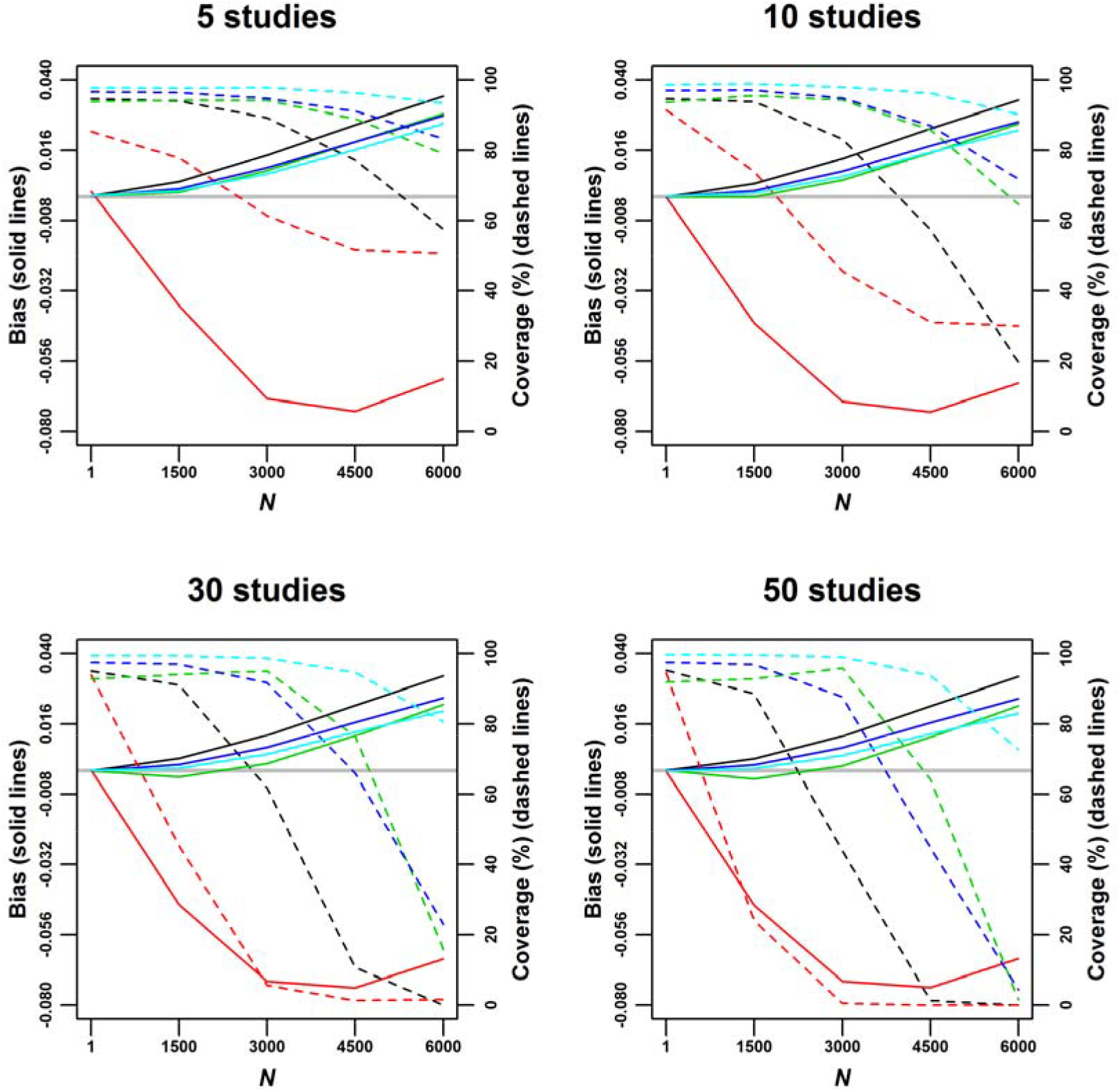
Bias (solid lines) and coverage (dashed lines) of the fixed effects (black), Egger regression (red), trim-and-fill (green), weighted median (dark blue) and mode-based estimate (light blue) under scenario 3: zero true effect (i.e., *β* = 0), small study effects through dissemination bias (assuming a linear relationship between *p*_*j*_ and *n*_*j*_), and study sizes uniformly ranging from 100 to 5000 individuals. *p*_*j*_: maximum P-value allowed for publication for a study with *n*_*j*_ participants. *N*: study size threshold, with studies larger than or equally sized to *N* not being affected by small study effects. The grey line indicates zero bias.

In scenario 4, small study effects resulted in a less marked asymmetry and in reduced *I*^2^ - i.e., under-dispersion (Supplementary Table 4). In general, the results were similar to scenario 3 (as shown in Supplementary Figure 2), with two main differences. First, the weighted median presented better coverage than the trim-and-fill, unless *K* = 50 and *N* = 4500. Second, the overcorrection presented by the trim-and-fill in scenario 3 was much more apparent, especially for larger values of *K*. Scenario 5 was in between scenarios 2 and 3 regarding *r* and *I*^2^ (Supplementary Table 5). In this scenario, the trim-and-fill was more biased than the weighted median and the MBE when the number of studies was low (*K* = 5 pr *K* = 10), and was in between them when there were more studies (*K* = 30 or *K* = 50). The difference between the weighted median and the MBE was small regardless of the number of studies (Supplementary Figure 3). In scenario 6, there was more between-study heterogeneity compared to the last scenario, but less than in scenario 2 (Supplementary Table 6). The weighted median and the MBE performed substantially better than the other methods (as shown in Supplementary Figure 4), with the MBE being close to unbiased in all cases when the number of studies was large (*K* = 30 or *K* = 50).

Supplementary Table 7 displays the performance of the methods to detect an effect in absence of small study effects (scenario 7). The fixed effects model was the most powered, followed by the trim-and-fill and the weighted median. Importantly, the trim-and-fill was slightly more precise than the weighted median, but presented substantially more power due to its under-coverage (which increased with number of studies and study size). The MBE was substantially more precise than Egger regression, but presented lower power due to under-coverage of the latter when the number of studies was low.

### 3.2. Real data examples

In our re-analysis of the catheter dataset (for which both *r* and *I*^2^ were high), the fixed effects model yielded an odds ratio of bloodstream infection of 0.47 (95% CI: 0.38; 0.58), while the weighted median and the MBE yielded the same smaller estimate of 0.57 (95% CI: 0.44; 0.75). Trim-and-fill yielded 0.45 (95% CI: 0.31; 0.65), similar to the fixed effects results. Egger regression yielded a qualitatively different estimate of 1.27 (95% CI: 0.70; 2.31) (Figure 5, panel A). This is likely an over-correction (as in the simulation study), especially given that the individual-study odds ratio estimates in the data ranged from 0.09 to 0.83.

**Figure 5.**
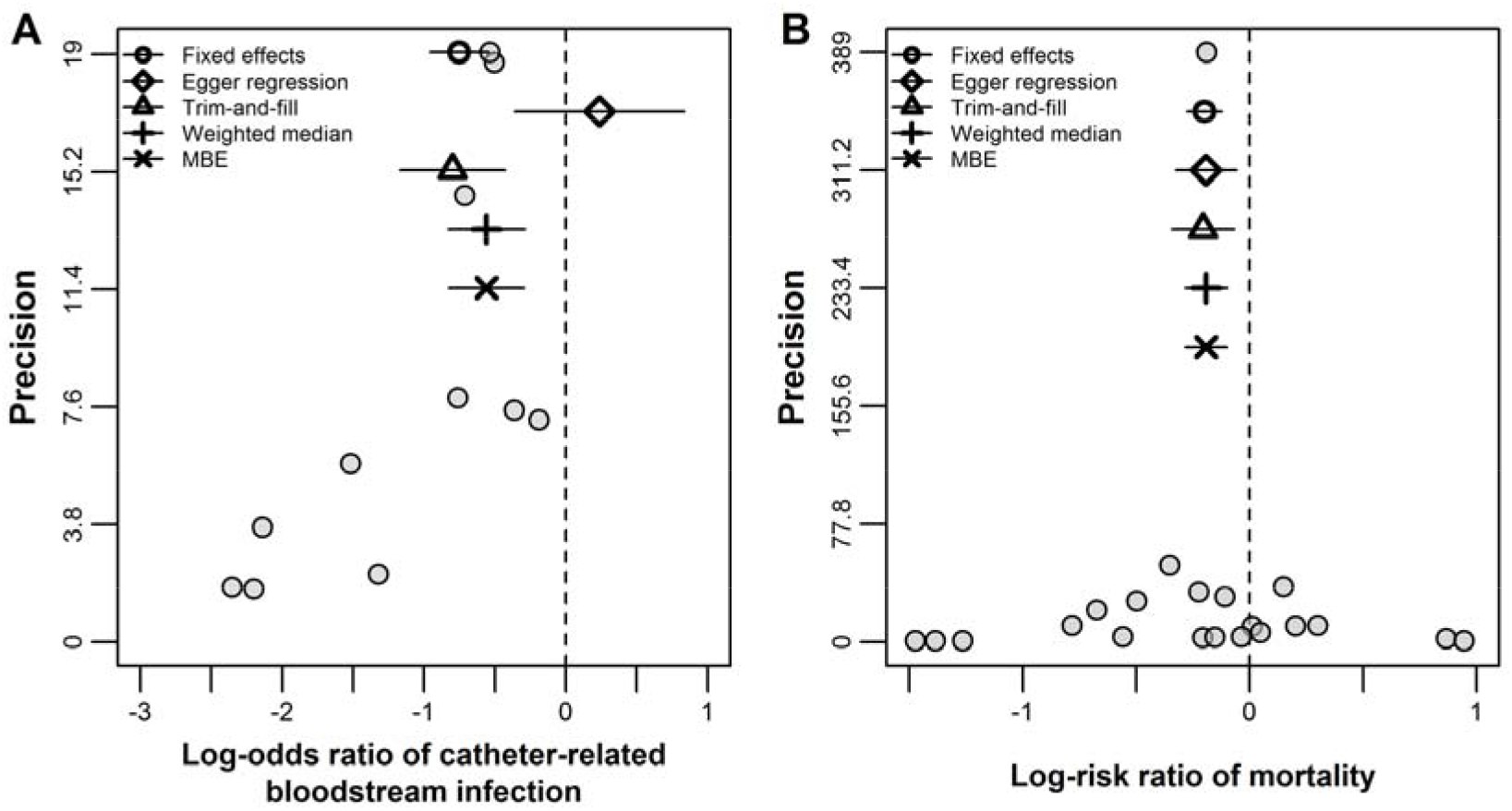
Funnel plots of the catheter (panel A) and streptokinase (panel B) meta-analyses, with pooled estimates and 95% confidence intervals from five meta-analysis methods. MBE: Mode-bases estimate.

For the aspirin dataset (which presented low *I*^2^ and marked asymmetry), the pooled odds ratio estimates of at least 50% of pain relief comparing active treatment to placebo were 3.43 (95% CI: 2.96; 3.98) for the fixed effect mean, 2.99 (95% CI: 2.41; 3.73) for the weighted median and 2.55 (95% CI: 1.78; 3.63) for the MBE. Trim-and-fill yielded an odds ratio of 2.87 (95% CI: 2.38; 3.47), which was in agreement with the weighted median and the MBE results. Egger regression yielded an odds ratio of 1.03 (95% CI: 0.71; 1.48), suggesting no effect of aspirin whatsoever (and again likely over-corrected).

For the streptokinase dataset (which was used as a positive control), the pooled risk ratio estimate comparing treatment and control groups was 0.82 (95% CI: 0.76; 0.88) for the fixed effects model. Results from the other four methods ranged from 0.81 to 0.83 (Figure 5, panel B). As a sensitivity analysis, the largest trial^13^ (which corresponded to a substantial proportion of the total weight in the meta-analysis) was removed, which had no material effect on the results.

## 4. Discussion

The results above suggest that the weighted median and MBE give sensible answers to real meta-analyses where small study effects are suspected (even when Egger regression or trim-and-fill do not), as well as similar results to the fixed effects model and other meta-analysis methods in absence of bias. This corroborates the results from the simulation study, which indicated that these methods are less influenced by small study effects than the conventional fixed effects model and other established methods (see the Supplementary Text for a discussion on bias due to small study effects in Egger regression). Software for their implementation is provided in the Supplementary Material.

There are several strategies to investigate the presence and degree of small study effects in meta-analysis, all of which have limitations.^14,15^ If, after careful examination, small study effects are suspected, we recommend that investigators apply the weighted median and the MBE in addition to standard methods as a sensitivity analyses. Our proposed approaches naturally reduce the influence of small studies without having to formally exclude them from the analysis. Exclusion often involves arbitrary study size cut-offs and artificially reduces the heterogeneity in the data.

When applying the weighted median and the MBE, it is important to not rely entirely on “statistical significance”, especially given that they will deliver estimates with less precision than the fixed effects model, but to examine their confidence intervals and assess their degree of overlap with the standard analysis. As a general rule, the weighted median and the MBE will give accurate and robust results when the majority of the weight in the analysis stems from studies that provide consistent effect estimates. This might, for example, be satisfied by just one or two large studies in a meta-analysis, despite the inclusion of many other biased studies. Conversely, they will give misleading results when the majority of the weight in the analysis stems from biased studies and, in the case of the MBE, the magnitude of the individual study biases are very similar (as illustrated in Figure 1, Panel D). The Cochrane Collaboration’s tool for assessing risk of bias^16^ could be used as a guide to the likely proportion of biased studies in a given meta-analysis, and to the value of applying of these techniques. As such the proposed methodology is a natural extension of exploring between study heterogeneity due to perceived risk of bias,^16^ which likely suffers from between rater subjectivity.

Importantly, the methods proposed here do not “correct” for asymmetry or heterogeneity between studies. Indeed, heterogeneity between studies should be expected,^17^ and exploring if measured study characteristics account for the latter (e.g., via subgroup analyses and meta-regression) may yield important insights regarding treatment effect modification and/or potential sources of bias. This cannot be achieved by simply applying the proposed methods, nor any other method that yields a single pooled point estimate. This is especially relevant for the MBE method, which assumes that there is a subset of homogeneous studies that yield consistent estimates of the treatment effect. Therefore, ideally the proposed methods would be applied if plausible effect modifiers do not account for observed heterogeneity between studies, or if there is residual heterogeneity within subgroups (although in this case the number of studies per subgroup may be prohibitive for meaningful comparisons between different estimators). Otherwise, the MBE can be used as a sensitivity analysis and interpreted as a test of the sharp null hypothesis; and, if the treatment effect can be assumed to be monotonic, as a test of the direction of the treatment effect. Finally, there are other waits to exploit the ZEMBA assumption (for example, by a model averaging approach^18^), and comparing their performance in plausible meta-analysis settings remains to be done.

Given its importance in summarising and interpreting the totality of available evidence, and their low cost compared to conducting a new study involving data collection, meta-analyses are particularly appealing to both researchers and journals, and a cornerstone of evidence-based medicine. Unfortunately, many systematic reviews and meta-analysis contain studies that are methodologically flawed and likely biased.^19^ We are confident that the weighted median and MBE provide inferences that are robust to small study effects under a variety of reasonable simulation models and in real datasets likely affected by this bias. We hope that these simple and intuitive methods will be used to strengthen the conclusions of meta-analyses.

