## Supplementary Materials for "The median and the mode as robust meta-analysis methods in the presence of small study effects"

**Supplementary Material**

Summary

**Supplementary Text**

#### Bias in Egger regression due to small study effects

In the catheter and aspirin meta-analyses (both of which presented evidence of funnel plot asymmetry), the pooled effect estimate from Egger regression ($\hat{\beta}_{Egger}$) was outside the range of individual study estimates (the $\hat{\beta}_{j}$’s, following the notation in equation (1)). More specifically, the catheter dataset presented a positive correlation between the $\hat{\beta}_{j}$’s and $\sigma_{j}^{-1}$’s ($r$=0.76), and its Egger regression pooled ln(odds ratio) estimate $\hat{\beta}_{Egger}$=0.24 was largest than $max(\hat{\beta}_{j})$=-0.19. The aspirin dataset presented a negative correlation between the $\hat{\beta}_{j}$’s and $\sigma_{j}^{-1}$’s ($r$=-0.70), and its Egger regression pooled ln(odds ratio) estimate $\hat{\beta}_{Egger}$=0.03 was smaller than $min(\hat{\beta}_{j})$=0.14. These results suggest, but do not prove, that Egger regression was overcorrecting for small study effects. This was corroborated in our simulations, where Egger regression yielded negative pooled estimates in the presence of positive bias and no treatment effect.

As in the main text, for clarity only we will momentarily assume that $b_{j}$ is the sole source of bias in equation (1) (i.e., only type (a) bias can occur). Egger regression therefore assumes a linear relationship between the $b_{j}$’s and $\sigma_{j}$’s, so that $b_{j}=\beta_{0}\sigma_{j}$. By plugging this expression for $b_{j}$ in equation (1), a simple manipulation leads to the Egger regression model:

$$\frac{\hat{\beta}_{j}}{\sigma_{j}}=\beta_{0}+\frac{\beta}{\sigma_{j}}+\varepsilon_{j} \left( \text{S1} \right)\text{.}$$

In equation (S1), $\beta_{0}$ is an intercept parameter that allows accounting for bias via non-zero $b_{j}$’s. For simplicity, equation (S1) shows the fixed effects Egger regression model, which can be extended into an additive or multiplicative random effects model;^1^ the latter was used in our simulations and real data examples.

Since Egger regression explicitly assumes that there is a linear relationship between the $b_{j}$’s and $\sigma_{j}$’s, it is prone to bias if the data-generating model leads to a non-linear relationship between the $b_{j}$’s and $\sigma_{j}$’s (i.e., if the Egger regression model is miss-specified).

Without restricting to type (a) bias, in our simulations bias can be defined as $b_{j}+{\sigma_{j}E[\varepsilon}_{j}|n_{j}]$, and the standard error as $\sigma_{j}\sqrt{\mathrm{Var}\left[ \varepsilon_{j}{|n}_{j} \right]}$. Therefore, the Egger regression model will only be correctly specified if $b_{j}+{\sigma_{j}E[\varepsilon}_{j}\left| n_{j} \right]=\beta_{0}\sigma_{j}\sqrt{\mathrm{Var}\left[ \varepsilon_{j}{|n}_{j} \right]}$; in other words, if $b_{j}+{\sigma_{j}E[\varepsilon}_{j}\left| n_{j} \right]$ and $\sigma_{j}\sqrt{\mathrm{Var}\left[ \varepsilon_{j}{|n}_{j} \right]}$ are linearly related. However, Supplementary Figure 5 shows that in all of our small study effects models the relationship between bias and standard error is non-linear, thus leading to bias in the Egger regression pooled effect estimate.

The small study effects models evaluated in our simulations represent plausible data-generating mechanisms that include the main features of typical models of small study effects. Particularly, in the case of dissemination bias, it is likely that selection is not influenced by precision itself, but by sample size (with larger studies being more likely to be published than smaller studies) and statistical significance levels (with studies that achieve conventional levels of statistical significance being more likely to be published than studies that do not), both of which are related to precision.

Our simulation results imply that Egger regression may suffer from bias in practice to the extent to which our small study effects models can be considered more or less plausible than the model assumed by Egger regression. If one assumes that all of these models are similarly plausible, our simulations then indicate that many plausible small study effects models may lead to substantial bias in Egger regression.

### Supplementary Tables

**Supplementary Table 1.** Performance of different meta-analysis methods under scenario 1: zero true effect (i.e., $\beta=0$), no small study effects, and study sizes uniformly ranging from 100 to 5000 individuals.

| **Method** | **Statistic** | $\boldsymbol{K}$ **(**$\boldsymbol{I}^{\boldsymbol{2}}$**;** $\boldsymbol{r}$**)** | | | |
| --- | --- | --- | --- | --- | --- |
|  |  | **5 (13.6%; 0.00)** | **10 (11.2%; 0.00)** | **30 (7.6%; 0.00)** | **50 (6.2%; 0.00)** |
| Fixed | Point estimate | 0.000 | 0.000 | 0.000 | 0.000 |
| Effects | Standard error | 0.018 | 0.013 | 0.007 | 0.006 |
|  | Coverage (%) | 95.2 | 94.9 | 94.9 | 95.4 |
|  | Power (%) | 4.8 | 5.1 | 5.1 | 4.6 |
| Egger | Point estimate | 0.000 | 0.000 | 0.000 | 0.000 |
| Regression | Standard error | 0.071 | 0.044 | 0.024 | 0.018 |
|  | Coverage (%) | 86.0 | 91.3 | 93.3 | 93.9 |
|  | Power (%) | 14.0 | 8.7 | 6.7 | 6.1 |
| Trim-and-fill | Point estimate | 0.000 | 0.000 | 0.000 | 0.000 |
|  | Standard error | 0.021 | 0.014 | 0.008 | 0.006 |
|  | Coverage (%) | 94.1 | 94.0 | 92.0 | 91.7 |
|  | Power (%) | 6.0 | 6.0 | 8.0 | 8.3 |
| Weighted | Point estimate | 0.000 | 0.000 | 0.000 | 0.000 |
| Median | Standard error | 0.022 | 0.017 | 0.010 | 0.008 |
|  | Coverage (%) | 96.8 | 97.2 | 97.6 | 97.7 |
|  | Power (%) | 3.2 | 2.8 | 2.4 | 2.3 |
| MBE | Point estimate | 0.000 | 0.000 | 0.000 | 0.000 |
|  | Standard error | 0.028 | 0.023 | 0.017 | 0.015 |
|  | Coverage (%) | 98.1 | 98.7 | 99.3 | 99.5 |
|  | Power (%) | 1.9 | 1.3 | 0.7 | 0.5 |

$K$: number of studies.

$I^{2}$: between-study heterogeneity.

$r$: Pearson correlation coefficient between point estimates ($\hat{\beta}_{j}$) and precision ($1/{\sigma_{j}}$).

MBE: mode-based estimate.

**Supplementary Table 2.** Between-study heterogeneity ($I^{2}$) and funnel plot asymmetry ($r$) according to the proportion of biased studies and number of studies ($K$) under scenario 2: zero true effect (i.e., $\beta=0$), small study effects through the bias term $b_{j}$, and study sizes uniformly ranging from 100 to 5000 individuals.

| **Statistic** | $\boldsymbol{K}$ | **Proportion (%) of biased studies (**$\boldsymbol{\delta}$ **parameter)** | | | | | | | | | | |
| --- | --- | --- | --- | --- | --- | --- | --- | --- | --- | --- | --- | --- |
|  |  | **0** | **10** | **20** | **30** | **40** | **50** | **60** | **70** | **80** | **90** | **100** |
| $I^{2}$ (%) | 5 | 13.5 | 38.7 | 56.7 | 69.7 | 78.2 | 83.1 | 86.3 | 87.2 | 87.3 | 85.9 | 83.5 |
|  | 10 | 11.3 | 49.9 | 70.2 | 80.7 | 86.0 | 88.8 | 89.7 | 90.3 | 90.1 | 89.5 | 88.0 |
|  | 30 | 7.9 | 64.5 | 80.3 | 86.1 | 88.6 | 90.0 | 90.7 | 91.0 | 90.9 | 90.4 | 89.5 |
|  | 50 | 6.4 | 68.1 | 81.7 | 86.6 | 88.9 | 90.2 | 90.8 | 91.1 | 91.0 | 90.5 | 89.6 |
| $r$ | 5 | 0.01 | -0.04 | -0.11 | -0.18 | -0.26 | -0.33 | -0.43 | -0.53 | -0.63 | -0.77 | -0.92 |
|  | 10 | 0.00 | -0.06 | -0.15 | -0.23 | -0.30 | -0.37 | -0.46 | -0.55 | -0.64 | -0.77 | -0.92 |
|  | 30 | 0.00 | -0.11 | -0.19 | -0.26 | -0.33 | -0.40 | -0.47 | -0.55 | -0.65 | -0.76 | -0.92 |
|  | 50 | 0.00 | -0.12 | -0.20 | -0.27 | -0.34 | -0.40 | -0.47 | -0.55 | -0.65 | -0.76 | -0.92 |

$r$: Pearson correlation coefficient between point estimates ($\hat{\beta}_{j}$) and precision ($1/{\sigma_{j}}$).

**Supplementary Table 3.** Between-study heterogeneity ($I^{2}$) and funnel plot asymmetry ($r$) according to the minimum study size required for not being affected by dissemination bias ($N$) and number of studies ($K$) under scenario 3: zero true effect (i.e., $\beta=0$), small study effects through dissemination bias (assuming a linear relationship between $p_{j}$ and $n_{j}$), and study sizes uniformly ranging from 100 to 5000 individuals.

| **Statistic** | $\boldsymbol{K}$ | $\boldsymbol{N}$ | | | | |
| --- | --- | --- | --- | --- | --- | --- |
|  |  | **1** | **1500** | **3000** | **4500** | **6000** |
| $I^{2}$ (%) | 5 | 13.5 | 15.9 | 18.9 | 17.3 | 14.5 |
|  | 10 | 11.4 | 14.1 | 16.4 | 15.1 | 11.2 |
|  | 30 | 7.7 | 10.8 | 14.8 | 12.5 | 7.2 |
|  | 50 | 6.4 | 9.7 | 13.9 | 11.4 | 5.5 |
| $r$ | 5 | 0.01 | -0.43 | -0.68 | -0.77 | -0.80 |
|  | 10 | 0.00 | -0.52 | -0.73 | -0.81 | -0.83 |
|  | 30 | 0.00 | -0.57 | -0.75 | -0.81 | -0.82 |
|  | 50 | 0.00 | -0.58 | -0.75 | -0.81 | -0.82 |

$p_{j}$: maximum P-value allowed for publication for a study with $n_{j}$ participants. $r$: Pearson correlation coefficient between point estimates ($\hat{\beta}_{j}$) and precision ($1/{\sigma_{j}}$).

**Supplementary Table 4.** Between-study heterogeneity ($I^{2}$) and funnel plot asymmetry ($r$) according to the minimum study size required for not being affected by dissemination bias ($N$) and number of studies ($K$) under scenario 4: zero true effect (i.e., $\beta=0$), small study effects through dissemination bias (assuming a square root relationship between $p_{j}$ and $n_{j}$), and study sizes uniformly ranging from 100 to 5000 individuals.

| **Statistic** | $\boldsymbol{K}$ | $\boldsymbol{N}$ | | | | |
| --- | --- | --- | --- | --- | --- | --- |
|  |  | **1** | **1500** | **3000** | **4500** | **6000** |
| $I^{2}$ (%) | 5 | 14.0 | 12.5 | 11.6 | 9.4 | 6.7 |
|  | 10 | 11.3 | 10.3 | 8.8 | 6.5 | 3.7 |
|  | 30 | 8.0 | 6.2 | 5.2 | 2.8 | 0.8 |
|  | 50 | 6.3 | 5.0 | 3.7 | 1.6 | 0.3 |
| $r$ | 5 | 0.00 | -0.32 | -0.52 | -0.62 | -0.64 |
|  | 10 | 0.00 | -0.39 | -0.58 | -0.66 | -0.69 |
|  | 30 | 0.00 | -0.43 | -0.61 | -0.69 | -0.71 |
|  | 50 | 0.00 | -0.44 | -0.62 | -0.69 | -0.71 |

$p_{j}$: maximum P-value allowed for publication for a study with $n_{j}$ participants. $r$: Pearson correlation coefficient between point estimates ($\hat{\beta}_{j}$) and precision ($1/{\sigma_{j}}$).

**Supplementary Table 5.** Between-study heterogeneity ($I^{2}$) and funnel plot asymmetry ($r$) according to the minimum study size required for not being affected by dissemination bias ($N$) and number of studies ($K$) under scenario 5: zero true effect (i.e., $\beta=0$), small study effects through dissemination bias (assuming a quadratic relationship between $p_{j}$ and $n_{j}$), and study sizes uniformly ranging from 100 to 5000 individuals.

| **Statistic** | $\boldsymbol{K}$ | $\boldsymbol{N}$ | | | | |
| --- | --- | --- | --- | --- | --- | --- |
|  |  | **1** | **1500** | **3000** | **4500** | **6000** |
| $I^{2}$ (%) | 5 | 13.3 | 26.0 | 36.1 | 37.0 | 34.0 |
|  | 10 | 11.5 | 25.4 | 38.6 | 40.4 | 34.9 |
|  | 30 | 7.6 | 25.8 | 41.6 | 43.5 | 37.9 |
|  | 50 | 6.5 | 26.1 | 42.5 | 44.7 | 39.2 |
| $r$ | 5 | 0.00 | -0.54 | -0.79 | -0.88 | -0.90 |
|  | 10 | 0.00 | -0.62 | -0.83 | -0.89 | -0.90 |
|  | 30 | 0.00 | -0.67 | -0.83 | -0.87 | -0.87 |
|  | 50 | 0.00 | -0.67 | -0.82 | -0.86 | -0.86 |

$p_{j}$: maximum P-value allowed for publication for a study with $n_{j}$ participants. $r$: Pearson correlation coefficient between point estimates ($\hat{\beta}_{j}$) and precision ($1/{\sigma_{j}}$).

**Supplementary Table 6.** Between-study heterogeneity ($I^{2}$) and funnel plot asymmetry ($r$) according to the $N_{small}$, $N_{large}$, and number of studies ($K$) under scenario 6: zero true effect (i.e., $\beta=0$), small study effects through dissemination bias (assuming a step function relationship between $p_{j}$ and $n_{j}$), and study sizes uniformly ranging from 100 to 5000 individuals.

| **Statistic** | $\boldsymbol{K}$ | $\boldsymbol{N}_{\boldsymbol{small}}\boldsymbol{;}\boldsymbol{N}_{\boldsymbol{large}}$ | | | |
| --- | --- | --- | --- | --- | --- |
|  |  | **0 ; 0** | **500 ; 1000** | **1000 ; 2000** | **2000 ; 4000** |
| $I^{2}$ (%) | 5 | 13.7 | 26.2 | 38.3 | 53.1 |
|  | 10 | 11.5 | 25.4 | 40.2 | 56.3 |
|  | 30 | 7.9 | 25.1 | 42.4 | 58.7 |
|  | 50 | 6.3 | 25.4 | 43.4 | 59.1 |
| $r$ | 5 | 0.00 | -0.27 | -0.51 | -0.81 |
|  | 10 | 0.00 | -0.37 | -0.63 | -0.85 |
|  | 30 | 0.00 | -0.47 | -0.69 | -0.86 |
|  | 50 | 0.00 | -0.49 | -0.70 | -0.87 |

$p_{j}$: maximum P-value allowed for publication for a study with $n_{j}$ participants. $N_{small}$: maximum sample size for a study to be classified as small. $N_{large}$: minimum sample size for a study to be classified as large. $r$: Pearson correlation coefficient between point estimates ($\hat{\beta}_{j}$) and precision ($1/{\sigma_{j}}$).

**Supplementary Table 7.** Performance of different meta-analysis methods under scenario 7: true effect $\beta=0$.02, no small study effects, and study sizes uniformly ranging from $n_{1}$ to $n_{2}$.

| **Method** | **Statistic** | $\boldsymbol{K}$ **(**$\boldsymbol{I}^{\boldsymbol{2}}$**;** $\boldsymbol{r}$**)** | | | | | | | | |
| --- | --- | --- | --- | --- | --- | --- | --- | --- | --- | --- |
|  |  | **5 (13.3%; 0.00)** | **10 (11.4%; 0.00)** | **30 (7.7%; 0.00)** | **50 (6.4%; 0.00)** | **5 (13.6%; 0.00)** | | **10 (11.5%; 0.00)** | **30 (7.7%; 0.00)** | **50 (6.3%; 0.00)** |
|  |  | $\boldsymbol{n}_{\boldsymbol{1}}\boldsymbol{=100;}\boldsymbol{n}_{\boldsymbol{2}}\boldsymbol{=1000}$ | | | | | $\boldsymbol{n}_{\boldsymbol{1}}\boldsymbol{=100}\boldsymbol{0;}\boldsymbol{n}_{\boldsymbol{2}}\boldsymbol{=5000}$ | | | |
| Fixed | Point estimate | 0.021 | 0.020 | 0.020 | 0.020 | 0.020 | | 0.020 | 0.020 | 0.020 |
| Effects | Standard error | 0.039 | 0.027 | 0.016 | 0.012 | 0.017 | | 0.012 | 0.007 | 0.005 |
|  | Coverage (%) | 95.2 | 95.2 | 94.9 | 94.6 | 94.7 | | 95.1 | 94.5 | 95.2 |
|  | Power (%) | 8.6 | 11.5 | 25.7 | 38.6 | 22.9 | | 41.1 | 85.0 | 97.1 |
| Egger | Point estimate | 0.021 | 0.021 | 0.020 | 0.020 | 0.019 | | 0.020 | 0.020 | 0.020 |
| Regression | Standard error | 0.184 | 0.114 | 0.063 | 0.048 | 0.099 | | 0.062 | 0.034 | 0.026 |
|  | Coverage (%) | 85.6 | 91.6 | 93.8 | 94.4 | 85.7 | | 91.7 | 93.7 | 94.2 |
|  | Power (%) | 14.2 | 8.6 | 7.5 | 7.5 | 15.1 | | 9.7 | 10.3 | 13.3 |
| Trim-and-fill | Point estimate | 0.021 | 0.020 | 0.020 | 0.020 | 0.020 | | 0.020 | 0.020 | 0.020 |
|  | Standard error | 0.043 | 0.030 | 0.017 | 0.013 | 0.018 | | 0.013 | 0.007 | 0.006 |
|  | Coverage (%) | 94.3 | 93.5 | 89.9 | 88.6 | 93.7 | | 93.1 | 88.0 | 84.6 |
|  | Power (%) | 8.9 | 12.0 | 26.8 | 38.1 | 20.8 | | 37.2 | 75.4 | 88.6 |
| Weighted | Point estimate | 0.021 | 0.020 | 0.020 | 0.020 | 0.020 | | 0.020 | 0.020 | 0.020 |
| Median | Standard error | 0.048 | 0.036 | 0.022 | 0.017 | 0.021 | | 0.016 | 0.009 | 0.007 |
|  | Coverage (%) | 96.3 | 97.1 | 97.6 | 97.3 | 96.6 | | 97.2 | 97.3 | 97.6 |
|  | Power (%) | 5.5 | 6.0 | 11.6 | 18.1 | 14.6 | | 23.5 | 56.9 | 80.8 |
| MBE | Point estimate | 0.020 | 0.021 | 0.021 | 0.020 | 0.020 | | 0.020 | 0.020 | 0.020 |
|  | Standard error | 0.063 | 0.053 | 0.040 | 0.035 | 0.028 | | 0.024 | 0.018 | 0.016 |
|  | Coverage (%) | 98.3 | 99.0 | 99.6 | 99.7 | 98.5 | | 99.2 | 99.7 | 99.8 |
|  | Power (%) | 2.9 | 2.0 | 2.5 | 2.8 | 6.6 | | 7.7 | 13.8 | 18.7 |

$K$: number of studies.

$I^{2}$: between-study heterogeneity.

$r$: Pearson correlation coefficient between point estimates ($\hat{\beta}_{j}$) and precision ($1/{\sigma_{j}}$).

MBE: mode-based estimate.

### Supplementary Figures

**Supplementary Figure 1. Illustration of the functional relationships between** $\boldsymbol{p}_{\boldsymbol{j}}$ **and** $\boldsymbol{n}_{\boldsymbol{j}}$ **induced by different dissemination bias models.**

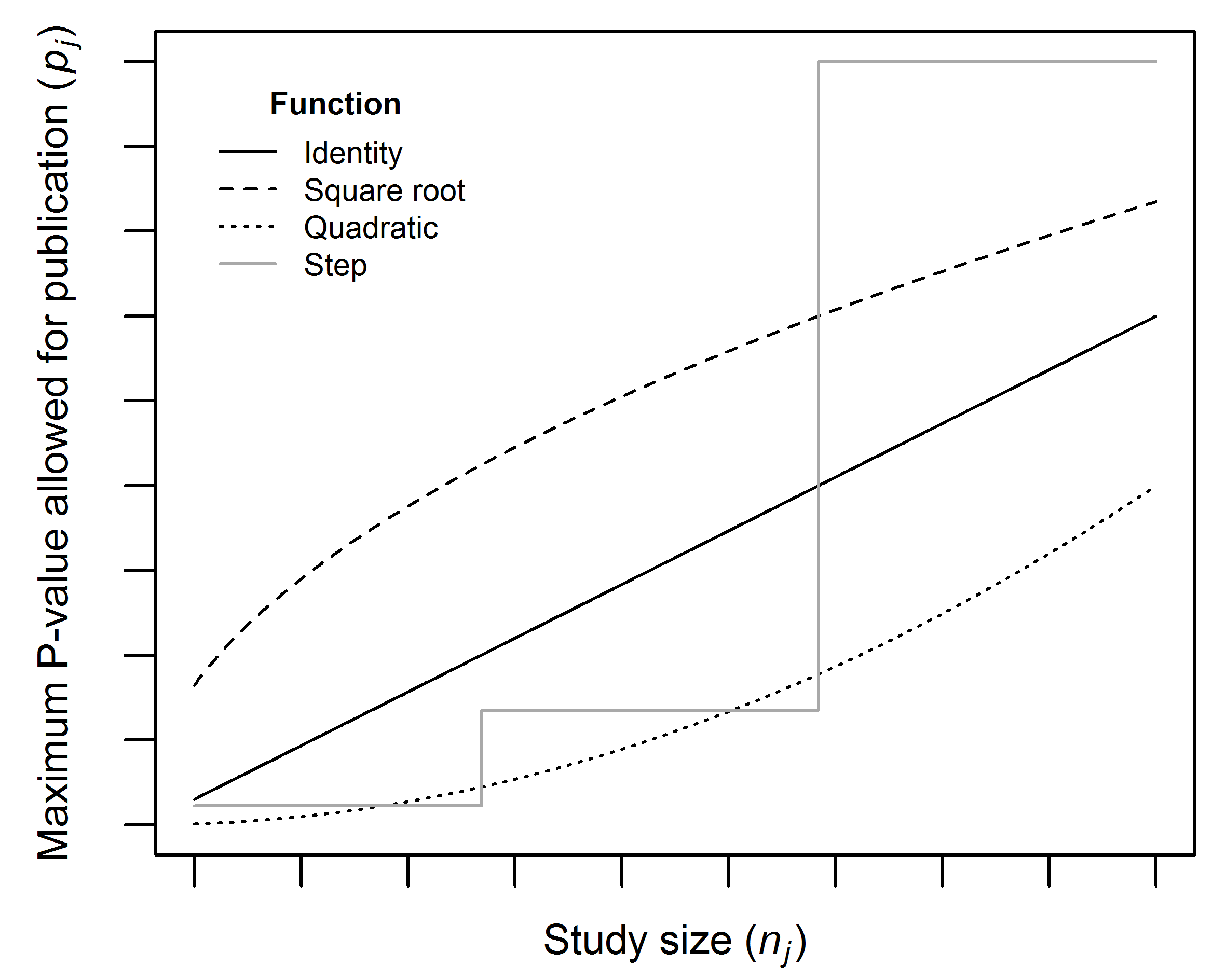

$p_{j}$: maximum P-value allowed for publication for a study with $n_{j}$ participants.

**Supplementary Figure 2. Bias (solid lines) and coverage (dashed lines) of the fixed effects (black), egger regression (red), trim-and-fill (green), weighted median (dark blue) and mode-based estimate (light blue) under scenario 4: zero true effect (i.e.,** $\boldsymbol{\beta=0}$**), small study effects through dissemination bias (assuming a square root relationship between** $\boldsymbol{p}_{\boldsymbol{j}}$ **and** $\boldsymbol{n}_{\boldsymbol{j}}$**), and study sizes uniformly ranging from 100 to 5000 individuals.**

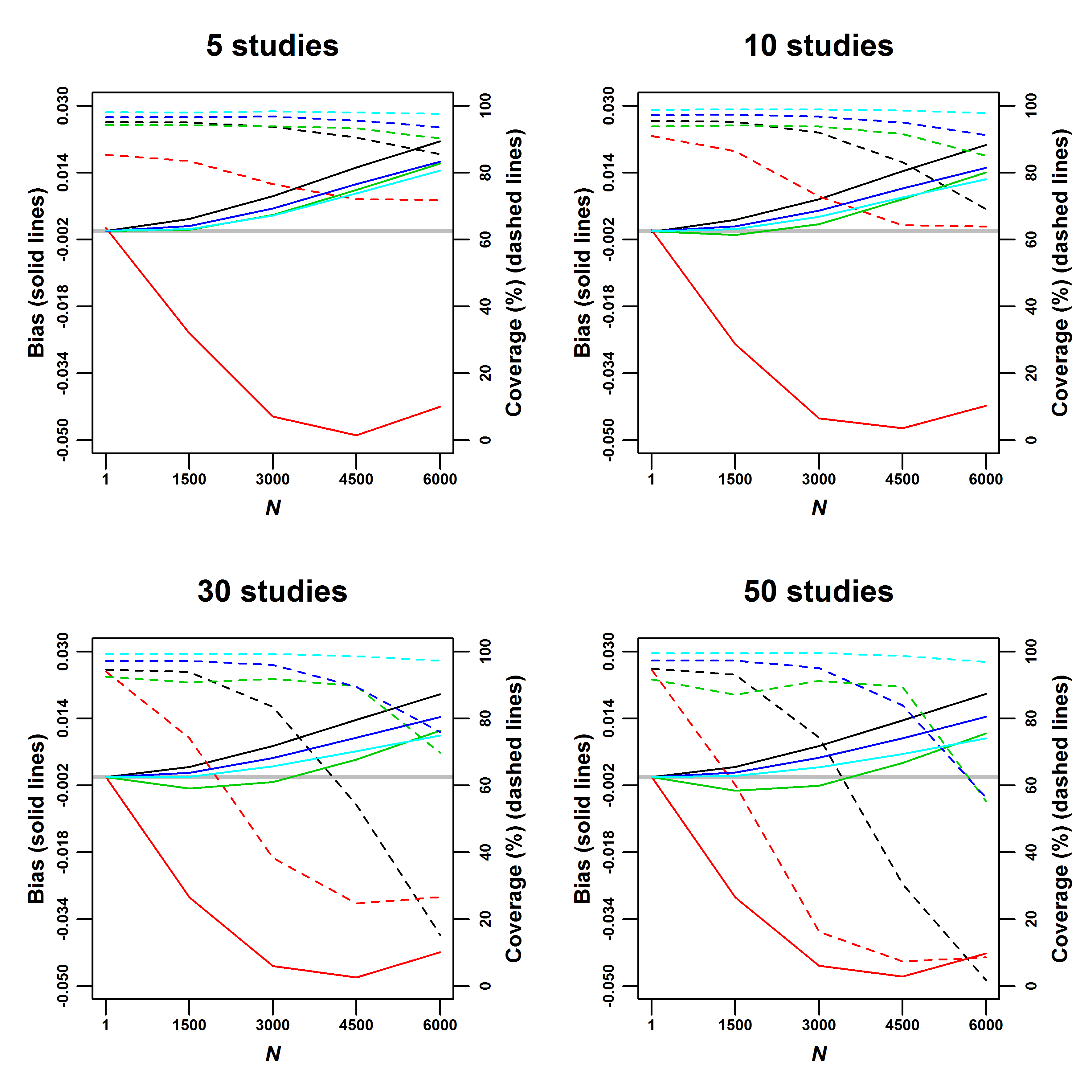

$p_{j}$: maximum P-value allowed for publication for a study with $n_{j}$ participants. $N$: study size threshold, with studies larger than or equally sized to $n^{*}$ not being affected by small study effects.

The grey line indicates zero bias.

**Supplementary Figure 3. Bias (solid lines) and coverage (dashed lines) of the fixed effects (black), egger regression (red), trim-and-fill (green), weighted median (dark blue) and mode-based estimate (light blue) under scenario 5: zero true effect (i.e.,** $\boldsymbol{\beta=0}$**), small study effects through dissemination bias (assuming a quadratic relationship between** $\boldsymbol{p}_{\boldsymbol{j}}$ **and** $\boldsymbol{n}_{\boldsymbol{j}}$**), and study sizes uniformly ranging from 100 to 5000 individuals.**

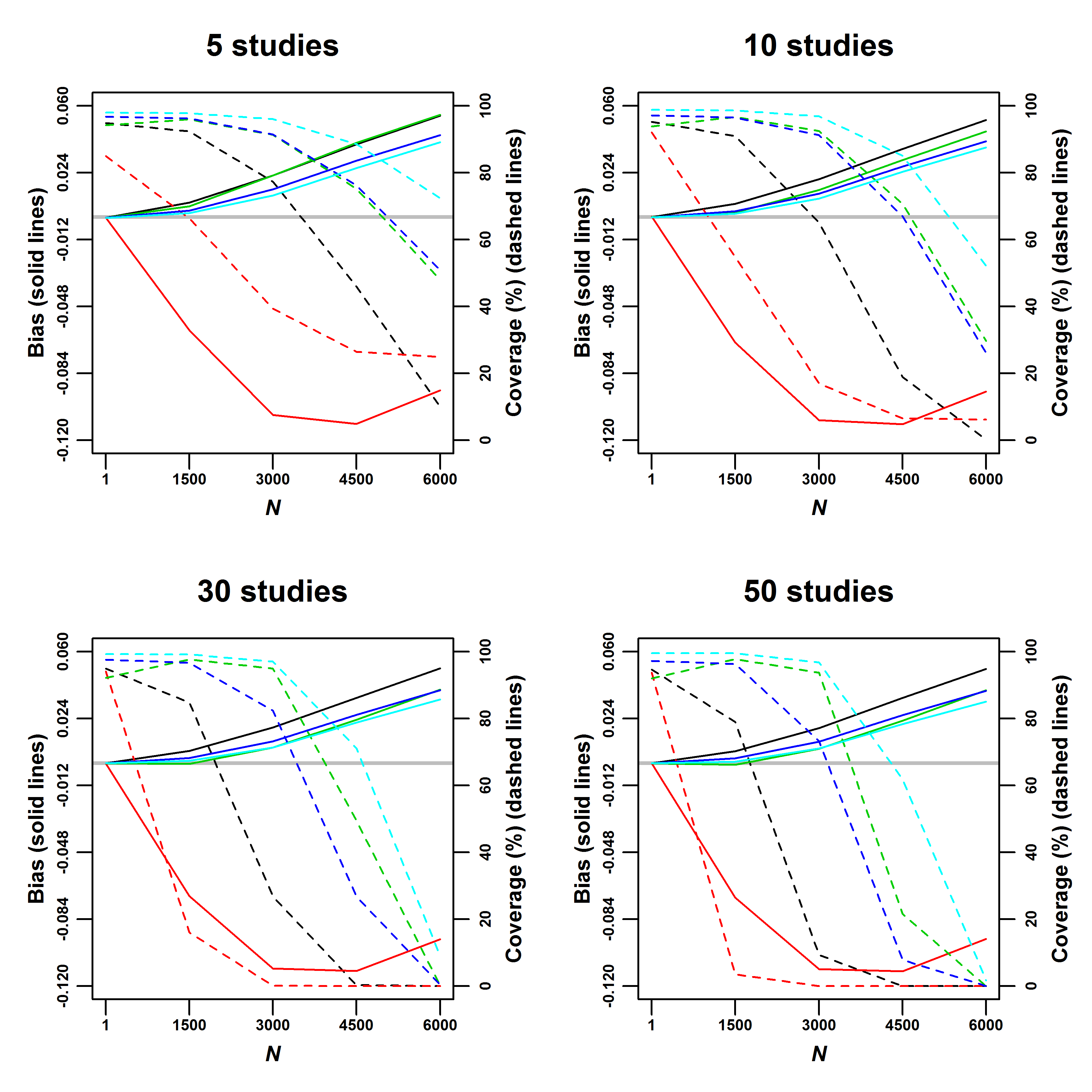

$p_{j}$: maximum P-value allowed for publication for a study with $n_{j}$ participants. $N$: study size threshold, with studies larger than or equally sized to $n^{*}$ not being affected by small study effects.

The grey line indicates zero bias.

**Supplementary Figure 4. Bias (solid lines) and coverage (dashed lines) of the fixed effects (black), egger regression (red), trim-and-fill (green), weighted median (dark blue) and mode-based estimate (light blue) under scenario 6: zero true effect (i.e.,** $\boldsymbol{\beta=0}$**), small study effects through dissemination bias (assuming a step function relationship between** $\boldsymbol{p}_{\boldsymbol{j}}$ **and** $\boldsymbol{n}_{\boldsymbol{j}}$**), and study sizes uniformly ranging from 100 to 5000 individuals.**

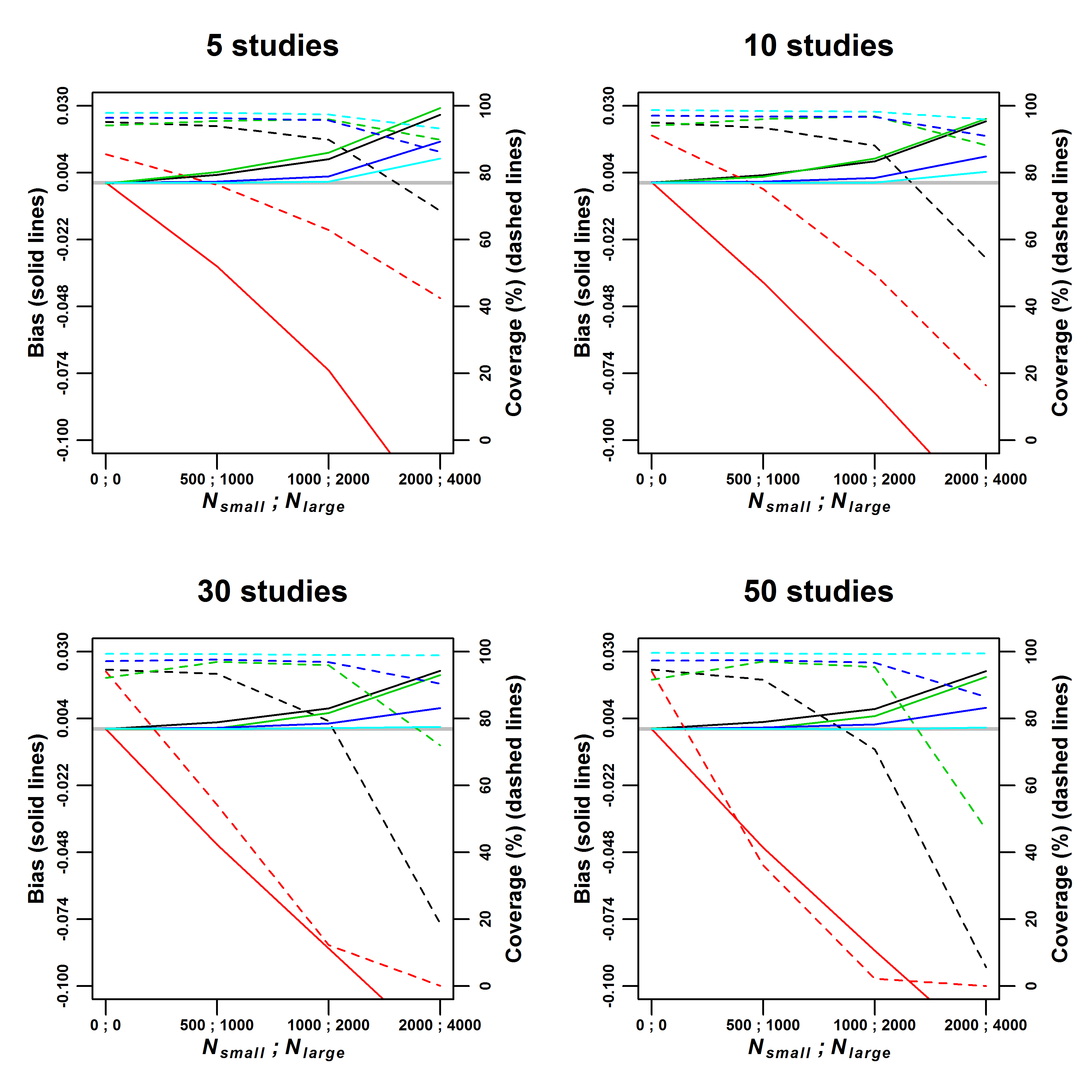

$p_{j}$: maximum P-value allowed for publication for a study with $n_{j}$ participants. $N_{small}$: minimum sample size for a study to be classified as medium-sized. $N_{large}$: minimum sample size for a study to be classified as large.

The grey line indicates zero bias.

**Supplementary Figure 5. Illustration of the relationship between bias and standard error induced by different models of small study effects.**

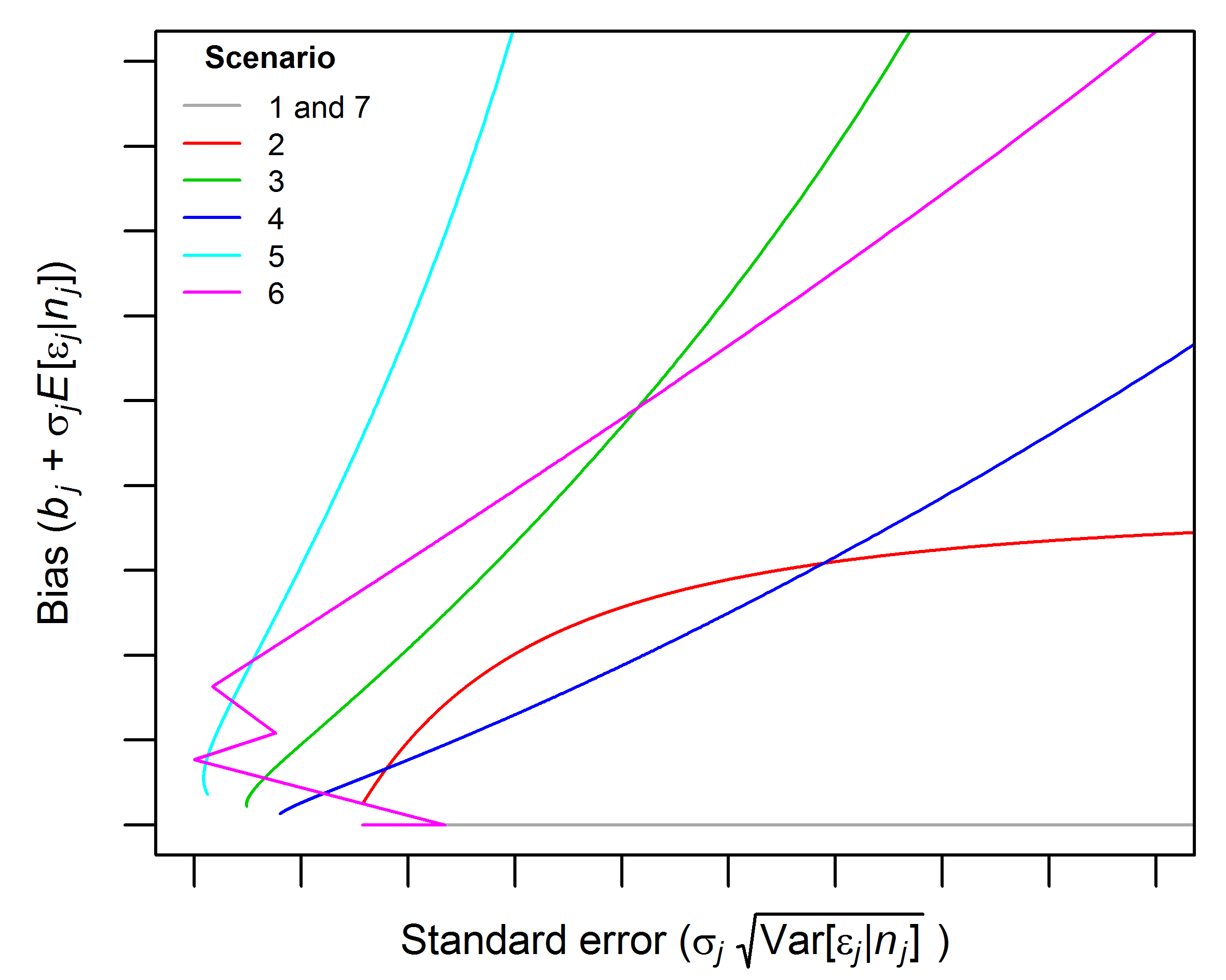

### Software code (R language)

#### Weighted median

```
#beta.in: point estimates (mean differences, log-odds ratio, etc.).
#se.in: standard errors
#alpha: confidence level of the confidence intervals. Defaults to 0.05 (i.e., 95% confidence intervals)
#n_boot: number of bootstrap iterations. Defaults to 1e4 (i.e., 10,000 iterations).

WeightedMedianMeta <- function(beta.in, se.in, alpha=0.05, n_boot=1e4) {

  #Function to calculate the point estimate
  weighted.median <- function(beta.in, weights.in) {
    beta.order          <- beta.in[order(beta.in)]
    weights.order       <- weights.in[order(beta.in)]
    weights.sum         <- cumsum(weights.order)-0.5*weights.order
    weights.sum         <- weights.sum/sum(weights.order)
    below               <- max(which(weights.sum<0.5))
    weighted.median.est <- beta.order[below] + (beta.order[below+1]-beta.order[below])*
      (0.5-weights.sum[below])/(weights.sum[below+1]-weights.sum[below])
    return(weighted.median.est)
  }

  #Calculate point estimate
  weights     <- se.in^-2                          #Inverse-variance weights
  pooled.beta <- weighted.median(beta.in, weights) #Inverse-variance weighted median

  #Calculate standard errors through bootstrapping
  boot.dist <- numeric(n_boot)
  for(a in 1:n_boot) {
    beta.boot     <- rnorm(n=length(beta.in), mean=beta.in, sd=se.in)
    boot.dist [a] <- weighted.median(beta.boot, weights)
  }
  pooled.se <- mad(boot.dist)

  #Calculate confidence intervals
  ci <- pooled.beta+c(-1,1)*qnorm(1-alpha/2)*pooled.se

  #Calculate P-value
  P <- pnorm(abs(pooled.beta)/pooled.se, lower.tail=F)*2

  #Provide results
  results        <- c(pooled.beta, pooled.se, ci, P)
  names(results) <- c('Beta', 'SE', 'CIlow', 'CIupp', 'Pvalue')
  return(results)
}
```

#### Mode-based estimate

```
#beta.in: point estimates (mean differences, log-odds ratio, etc.).
#se.in: standard errors
#alpha: confidence level of the confidence intervals. Defaults to 0.05 (i.e., 95% confidence intervals)
#n_boot: number of bootstrap iterations. Defaults to 1e4 (i.e., 10,000 iterations).

MBEMeta <- function(beta.in, se.in, alpha=0.05, n_boot=1e4) {

  #Function to calculate the point estimate
  MBE <- function(beta.in, weights.in) {
    bw       <- 0.9*(min(sd(beta.in), mad(beta.in)))/length(beta.in)^(1/5) #Bandwidth
    weights  <- weights.in/sum(weights.in)                                 #Standardising weights
    EDF      <- density(beta.in, bw=bw, weights=weights)                   #Weighted empirical density function
    MBE.est <- EDF$x[which.max(EDF$y)]                                     #Calculate point estimate
    return(MBE.est)
  }

  #Calculate point estimate
  weights     <- se.in^-2              #Inverse-variance weights
  pooled.beta <- MBE(beta.in, weights) #Inverse-variance weighted MBE

  #Calculate standard errors through bootstrapping
  boot.dist <- numeric(n_boot)
  for(a in 1:n_boot) {
    beta.boot   <- rnorm(n=length(beta.in), mean=beta.in, sd=se.in)
    boot.dist [a] <- MBE(beta.boot, weights)
  }
  pooled.se <- mad(boot.dist)

  #Calculate confidence intervals
  ci <- pooled.beta+c(-1,1)*qnorm(1-alpha/2)*pooled.se

  #Calculate P-value
  P <- pnorm(abs(pooled.beta)/pooled.se, lower.tail=F)*2

  #Provide results
  results        <- c(pooled.beta, pooled.se, ci, P)
  names(results) <- c('Beta', 'SE', 'CIlow', 'CIupp', 'Pvalue')
  return(results)
}
```
